# Emergence of IS*Aba1*-linked oxacillinase genes among carbapenem resistant *Acinetobacter baumannii* isolates in a tertiary cardiac center, Nepal

**DOI:** 10.1101/2023.03.13.532405

**Authors:** Shrijana Bista, Bindeshwar Yadav, Gopiram Syangtan, Jivan Shakya, Reshma Tuladhar, Dev Raj Joshi, Binod Lekhak

## Abstract

Insertion sequence contributes to the emergence of carbapenem resistance by dissemination of carbapenemase genes and providing promoter for their overexpression. This study aims to ascertain the occurrence of IS*Aba1*-linked OXA carbapenemase genes and its relevance to carbapenem resistance level in *Acinetobacter baumannii*. This hospital based descriptive study was conducted at Shahid Gangalal National Heart Center, Kathmandu, Nepal. An overall of 1,291 clinical specimens received for routine culture and antibiotic susceptibility testing throughout the study period were included in this study. Identification of *Acinetobacter baumannii* was validated through detection of intrinsic *bla*_OXA-51-like_ gene by polymerase chain reaction (PCR). Antibiotic susceptibility was tested by Kirby-Bauer disc diffusion approach and minimum inhibitory concentration (MIC) of meropenem was assessed through agar dilution method. Uniplex PCR assays were performed to detect genes encoding oxacillinases and IS*Aba1*. Upstream association of insertion element, IS*Aba1* to oxacillinase genes was assessed through PCR mapping strategy using IS*Aba1*F and OXA-51R/OXA-23R primers. Out of the 340 bacteria isolated, only 40 (11.8%) were *Acinetobacter baumannii*. All isolates were resistant against meropenem with MIC value ranging from 16-256 μg/ml. *bla*_OXA-23-like_ genes was present in every isolate but *bla*_OXA-58_ in just two isolates (5%). All isolates had IS*Aba1* either above *bla*_OXA-23-like_ or *bla*_OXA-51-like_ gene. Higher MIC_90_ value of meropenem (243.20 μg/ml) was found in *A*. *baumannii* cluster with IS*Aba1*-linked upstream to both *bla*_OXA-23-like_ and *bla*_OXA-51-like_ genes, thus depicting their eminent role to enhanced carbapenem resistance. *Acinetobacter baumannii* isolates with IS*Aba1*-linked oxacillinase genes are rapidly emerging in clinical settings of Nepal. Thus, medical communities need to be prepared and enable targeted approaches for managing burgeoning problem of carbapenem resistance.

## Introduction

*Acinetobacter baumannii* has become an obstinate nosocomial bacteria [1], in virtue of its multi-resistance phenotype [2], especially among the immune-compromised patients and mostly in intensive care units (ICUs) [3]. Due to their high genetic plasticity, they are able to acquire different resistance determinants through horizontal gene transfer by mobile genetic elements (MGEs) [4]. Despite the fact that carbapenems are the preferred medication to treat infections caused by multidrug resistant (MDR) *Acinetobacter baumannii*, resistance against this antibiotic is escalating at an alarming rate throughout the world [5]. The World health organization (WHO) has also listed carbapenem resistant *A*. *baumannii* to be one of the 12 critical priority pathogens for research of new antibiotics [6]. The underlying resistance mechanisms includes overexpression of efflux pumps, modifications to outer membrane proteins and carbapenemases encoded even on plasmids, integrons and transposons [7,8].

The carbapenem resistance phenotype in *Acinetobacter baumannii* is chiefly due to the acquisition of oxacillinase genes flanked by insertion sequence (IS) elements [9]. Insertion sequence employs a crucial part in the propagation and expression of antibiotic resistance genes in *A*. *baumannii* [10]. They provide promoter for overexpression of various downstream antibiotic resistance genes [11,12]. Despite the significant prevalence of carbapenem resistance in *A*. *baumannii* isolates [13,14], there is paucity of studies focused to unravel the prevailing carbapenem resistance mechanism in Nepal. Therefore, this study was carried out in order to investigate carbapenem resistance profile, occurence of OXA carbapenemase genes (*bla*_OXA-51-like_, *bla*_OXA-23-like_, *bla*_OXA-58_), insertion sequence (IS*Aba1*), and IS*Aba1*-linked oxacillinase genes and their relationship to carbapenem susceptibility among *A*. *baumannii* isolates from cardiac patients in Nepal.

## Materials and methods

### Clinical settings and study design

This hospital based cross-sectional study was conducted from April 2021 to March 2022 at Shahid Gangalal National Heart Center (SGNHC), a tertiary referral cardiac center in, Kathmandu, Nepal. The study population included patients visiting to SGNHC irrespective of any age and gender within the study period. In the study, all clinical samples that were received for routine culture and testing for antibiotic susceptibility were included. A total of 1,291 clinical specimens were processed and analyzed consistent with standard microbiological practices [15] during the study period.

Initially, isolates were classified as *Acinetobacter calcoaceticus* – *Acinetobacter baumannii* (ACB) complex utilizing cultural, biochemical and microscopic methods like growth on MacConkey agar, gram staining, oxidase, motility, oxidation-fermentation (OF), citrate utilization, triple sugar iron agar (TSIA) and growth at 42°C [16]. After that, they were delivered to Tribhuvan University’s Central Department of Microbiology where they were kept at −20°C in tryptic soy broth that had been added with 20% glycerol (v/v) until a subsequent downstream investigation. ACB complex were further characterized as *Acinetobacter baumannii* by observation of intrinsic *bla*_OXA-51-like_ gene [17]. As the reference strain, *A*. *baumannii* ATCC 19606 was employed.

### Antibiotic susceptibility testing (AST)

*Acinetobacter baumannii* isolates were subjected to in-vitro antibiotic susceptibility towards 16 clinically relevant antibiotics through modified Kirby-Bauer disc diffusion technique. Different antibiotics tested were ampicillin/sulbactam (10/10 μg), piperacillin/tazobactam (100/10 μg), cefepime (30 μg), cefotaxime (30 μg), ceftazidime (30 μg), ciprofloxacin (5 μg), meropenem (10 μg), imipenem (10 μg), gentamicin (10 μg), amikacin (30 μg), tigecycline (15 μg), tetracycline (30 μg), doxycycline (30 μg), polymyxin B (300 U), colistin (10 μg) and cotrimoxazole (25 μg). Results were analyzed in accordance with the susceptibility breakpoints established by the Clinical and Laboratory Standards Institute (CLSI) [18]. The interpretation of tigecycline, colistin, and polymyxin B susceptibility was based on breakpoints as outlined by Galani [19], Kwa [20] and Jones [21] respectively. As a quality control measure, antibiotic susceptibility assays were validated using *Escherichia coli* ATCC 25922 and *Pseudomonas aeruginosa* ATCC 27853 as reference strains.

Multi Drug Resistant (MDR) is a designation for isolates that are resistant to at least one agent in three or more antimicrobial categories, while Extensively Drug Resistant (XDR) is used to describe *Acinetobacter baumannii* isolates that are only susceptible to one or two categories [22].

Agar dilution technique was used to evaluate the MIC of meropenem (Sigma Aldrich, USA) against all isolates of *Acinetobacter baumannii*. The findings were interpreted in accordance with the clinical breakpoint of (Susceptible ≤ 2 and Resistant ≥ 8 μg/ml) as recommended by CLSI [18].

### Phenotypic detection of β-lactamase producers

Combined disc test (CDT) was used to phenotypically identify the presence of various β-lactamases, including ESBL (Extended spectrum beta lactamase) [18], MBL (Metallo beta lactamase) [23], KPC (*Klebsiella pneumoniae* carbapenemase) [24], and AmpC (Ampicillinase C) [25].

### Detection of oxacillinases and IS*Aba1* genes

Total genomic DNA was extracted by heat shock method as stated by Hartas [26]. DNA extracted, including its quantity and quality was evaluated by spectrophotometric analysis using (Nanodrop 1000, Thermofischer Scientific, USA).

The Uniplex PCR assays were conducted to assess the presence of genes encoding oxacillinases, including *bla*_OXA-51-like_, *bla*_OXA-23-like_, *bla*_OXA-58_ as well as IS*Aba1* genes using specific primers as described previously [27–30]. All primers were purchased from Macrogen Inc. (South Korea) and the expected amplicon sizes and primer sequences are listed in (Table 1). PCR reactions were conducted in 25 μL reaction volumes with 12.5 μL of 2x master mix (Biolabs, USA), 1 μL of each forward and reverse primer, 3 μL of template DNA and 7.5 μL of nuclease free water. The gradient thermal cycler (Applied Biosystems, USA) was used for the amplification, and the target genes’ specific thermal cycling settings are specified in (Table S1). Nuclease-free water served as the negative control in each PCR cycle. *Acinetobacter baumannii* isolates previously confirmed to have *bla*_OXA-23-like_, *bla*_OXA-58_ and IS*Aba1* genes [10] were kindly provided by Professor Dr. Balaji Veeraraghavan, Department of Microbiology, Christian Medical College, Vellore, India and were used as positive amplification controls. The PCR products were separated by electrophoresis on a 1.5% (w/v) agarose gel prepared in tris acetic acid – EDTA (TAE) buffer with 0.5 μg/ml ethidium bromide and observed in gel documentation system (Azure Biosystems, USA). The amplicon sizes were determined by comparison with 100 bp DNA molecular size marker (GenScript, Singapore).

**Table 1:**
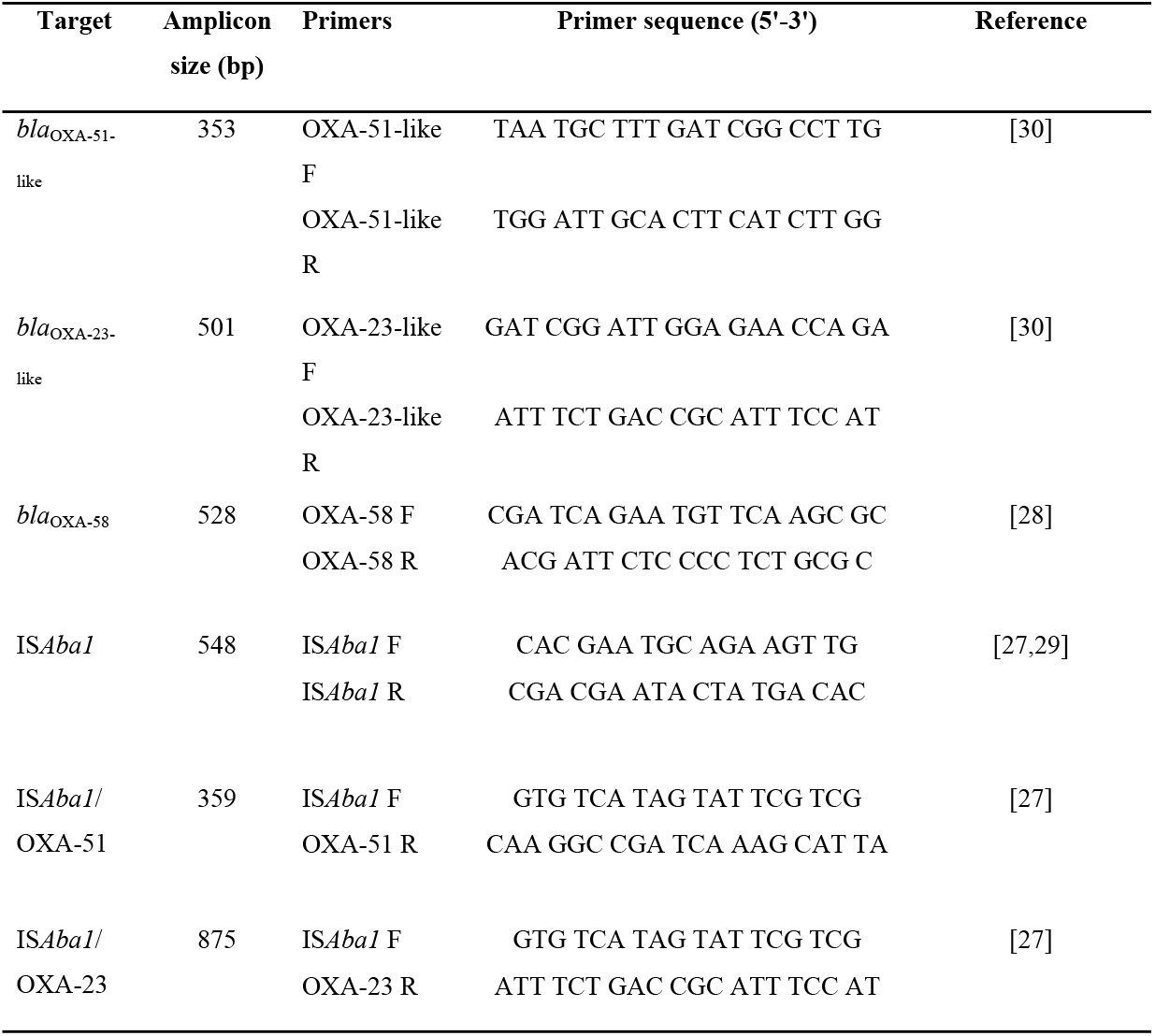
Specific primers used in this study for the amplification of target genes

### Detection of IS*Aba1* associated with OXA-51 and OXA-23

The position of the IS*Aba1* sequence in relation to the *bla_OXA_* genes was investigated with the polymerase chain reaction as described previously [27]. Upstream association of insertion sequence, IS*Aba1* to *bla*_OXA-51-like_ and *bla*_OXA-23-like_ gene was evaluated through PCR mapping approach using the primer pairs IS*Aba1*F/OXA-51R and IS*Aba1*F/OXA-23R respectively.

### Data analysis

All the data obtained were entered into the Excel 2013 (Microsoft Corp., Redmond, USA). Statistical Package for the Social Sciences software version 23 (SPSS, Inc. Chicago, USA) was employed for descriptive statistical assessment. Chi-square analysis was used to examine categorical data, with a p-value of less than 0.05 indicating statistical significance.

## Results

### Microbiological and demographic attributes

Out of the total 1,291 clinical specimens processed, 26.3% (340/1,291) showed bacterial growth. Among 340 culture positive isolates, *Acinetobacter* spp. accounted for (13.8%; 47/340). Of these, 45 (95.7%) were identified to be ACB complex and 2 isolates belonged to other *Acinetobacter* spp. on account of various biochemical tests and growth temperatures. Of the 45 ACB complex, 40 (88.9%) were identified to be *A*. *baumannii* based on the presence of an endogenous *bla*_OXA-51-like_ gene. The prevalence of *A*. *baumannii* among total bacteria isolated was thus 11.8%.

There was comparatively greater incidence of *Acinetobacter baumannii* infection in male (57.5%) than female patients (42.5%). The median age of patients infected from *A*. *baumannii* was 40 years (ranging from 1 month to 86 years of age). Patients under ten years old had maximum prevalence of *Acinetobacter baumannii* infection (n = 9, 22.5%).

All the 40 isolates of *Acinetobacter baumannii* were from inpatient source. Of them, predominant isolates (60%) came from ICU. The most prevalent specimen to yield *A*. *baumannii* isolates was sputum (57.5%) proceeded by endotracheal secretion (20%). Equal number of isolates (10% each) was recovered from blood and wound swab and least only 1 (2.5%) isolate was obtained from urine sample.

### Antibiotic susceptibility

Antibiotic susceptibility profiles through disc diffusion revealed that all the isolates were resistant to extended spectrum cephalosporins (cefepime, cefotaxime and ceftazidime), ciprofloxacin and piperacillin/tazobactam. The percentages of isolates that were resistant to the other antibiotics analyzed were as follows: Gentamicin (97.5%), meropenem (97.5%), imipenem (92.5%), amikacin (80%), cotrimoxazole (75%), ampicillin/sulbactam (65%), tetracycline (55%) and doxycycline (47.5%). The highest rate of susceptibility was observed against polymyxin B (100%), followed by tigecycline (80%) and colistin (77.5%). The isolates were all multidrug resistant (MDR), with 30% of them even being extremely drug resistant (XDR).

The *Acinetobacter baumannii* isolates were all found to be meropenem-resistant based on their MIC values. Meropenem’s MICs ranged between 16 and 256 μg/ml, with MIC_50_ and MIC_90_ values of 64 and 128 μg/ml respectively. A high MIC value of 64 μg/ml or higher was present in 72.5% of the isolates.

### β-lactamase profile of *Acinetobacter baumannii*

On the phenotypic β-lactamase production test by combined disc method, 30% of the *Acinetobacter baumannii* isolates were identified to be MBL producer. Two of the MBL producer also co-produced ESBL and KPC each, whereas 10% of the isolates only produced ESBL. Similarly, only one isolate (2.5%) produced AmpC β-lactamase enzyme. Moreover, there was significant association between β-lactamase profile and resistance phenotypes of *Acinetobacter baumannii* (p = 0.04). Every ESBL producing *A*. *baumannii* isolates were exclusively MDR, while more number of only MBL generating *A*. *baumannii* isolates was XDR (Table 2).

**Table 2:** Association of β-lactamase profile of *A*. *baumannii* with resistance phenotypes

| $\beta$ – lactamase profile | Resistance phenotypes | | Total | p – value* |
| --- | --- | --- | --- | --- |
|  | MDR<br>No. (%) | XDR<br>No. (%) |  |  |
| **AmpC** | 1 (2.5) | 0 (0) | 1 (2.5) |  |
| **ESBL only** | 4 (10) | 0 (0) | 4 (10) |  |
| **MBL only** | 3 (7.5) | 7 (17.5) | 10 (25) |  |
| **MBL + ESBL** | 1 (2.5) | 0 (0) | 1 (2.5) | **0.04** |
| **MBL + KPC** | 1 (2.5) | 0 (0) | 1 (2.5) |  |
| **Non-β-lactamase** | 18 (45) | 5 (12.5) | 23 (57.5) |  |
| **Total** | 28 (70) | 12 (30) | 40 (100) |  |

\*Chi-square test

### Molecular characterization of *Acinetobacter baumannii*

All 40 carbapenem resistant *Acinetobacter baumannii* isolates were discovered to co-harbor *bla*_OXA-51-like_, *bla*_OXA-23-like_ and insertion sequence, IS*Aba1* genes. Among them, 2 (5%) isolates also harbored *bla*_OXA-58_ gene in addition to these genes.

Similarly, all 40 (100%) of the *Acinetobacter baumannii* isolates had IS*Aba1* gene either linked to *bla*_OXA-23-like_ or resident *bla*_OXA-51-like_ genes. Of them, 30 (75%) isolates gave a band of 875 bp in a PCR utilizing the IS*Aba1*F and OXA-23R primer pairs, stating plausible upstream location of IS*Aba1* above *bla*_OXA-23-like_ gene. On the same line, IS*Aba1* was detected upstream of *bla*_OXA-51-like_ gene in 30 (75%) of the isolates, as a band of around 359 bp was observed in a PCR employing the IS*Aba1*F and OXA-51R primers. IS*Aba1* was solely found upstream of *bla*_OXA-51-like_ and *bla*_OXA-23-like_ genes in each 10 of the isolates, whereas IS*Aba1* lay upstream of both *bla*_OXA-51-like_ and *bla*_OXA-23-like_ genes in 20 (50%) of the *A*. *baumannii* isolates (Table 3).

**Table 3:**
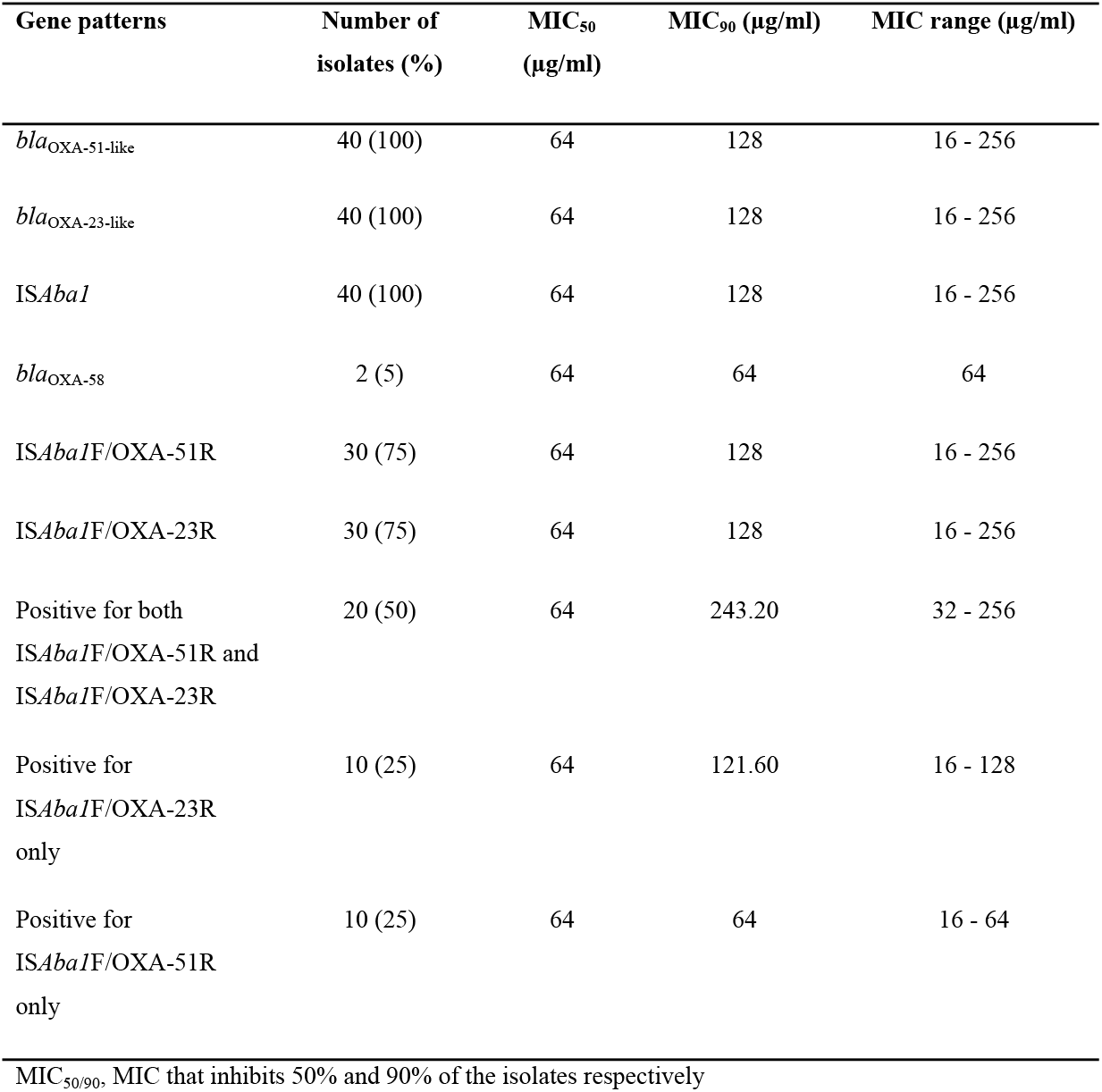
Meropenem resistance level in relation to distribution of IS*Aba1*, oxacillinase and IS*Aba1*-linked oxacillinase genes

### Relevance of meropenem resistance level to distribution of IS*Aba1*-linked oxacillinase genes

In this study, 20 *Acinetobacter baumannii* isolates in which IS*Aba1* had upstream association to both *bla*_OXA-51like_ and *bla*_OXA-23-like_ genes had comparatively higher MIC_90_ value of 243.20 μg/ml, than that each of the 10 isolates in which IS*Aba1* had upstream association only to *bla*_OXA-23-like_ gene (121.60 μg/ml) and *bla*_OXA-51-like_ gene (64 μg/ml) (Table 3). This was further illustrated in (Fig. 1), in which isolates positive for both IS*Aba1*F/OXA-51R and IS*Aba1*F/OXA-23R had comparatively higher MIC values for meropenem. Additionally, the highest MIC value of 256 μg/ml was only present in isolates with IS*Aba1* linked upstream of both *bla*_OXA-51-like_ and *bla*_OXA-23-like_ genes (Fig. 1). Moreover, *A*. *baumannii* isolates with IS*Aba1* upstream of *bla*_OXA-23-like_ gene regardless of the status of upstream insertion of IS*Aba1* to *bla*_OXA-51-like_ gene demonstrated higher MICs of meropenem (16-256 μg/ml) with MIC_90_ value of 128 μg/ml. Despite IS*Aba1* being upstream to *bla*_OXA-51-like_ gene, isolates without upstream insertion of IS*Aba1* to *bla*_OXA-23-like_ gene displayed relatively low MIC of meropenem (16-64 μg/ml), with only 64 μg/ml of MIC_90_ value.

**Figure 1:**
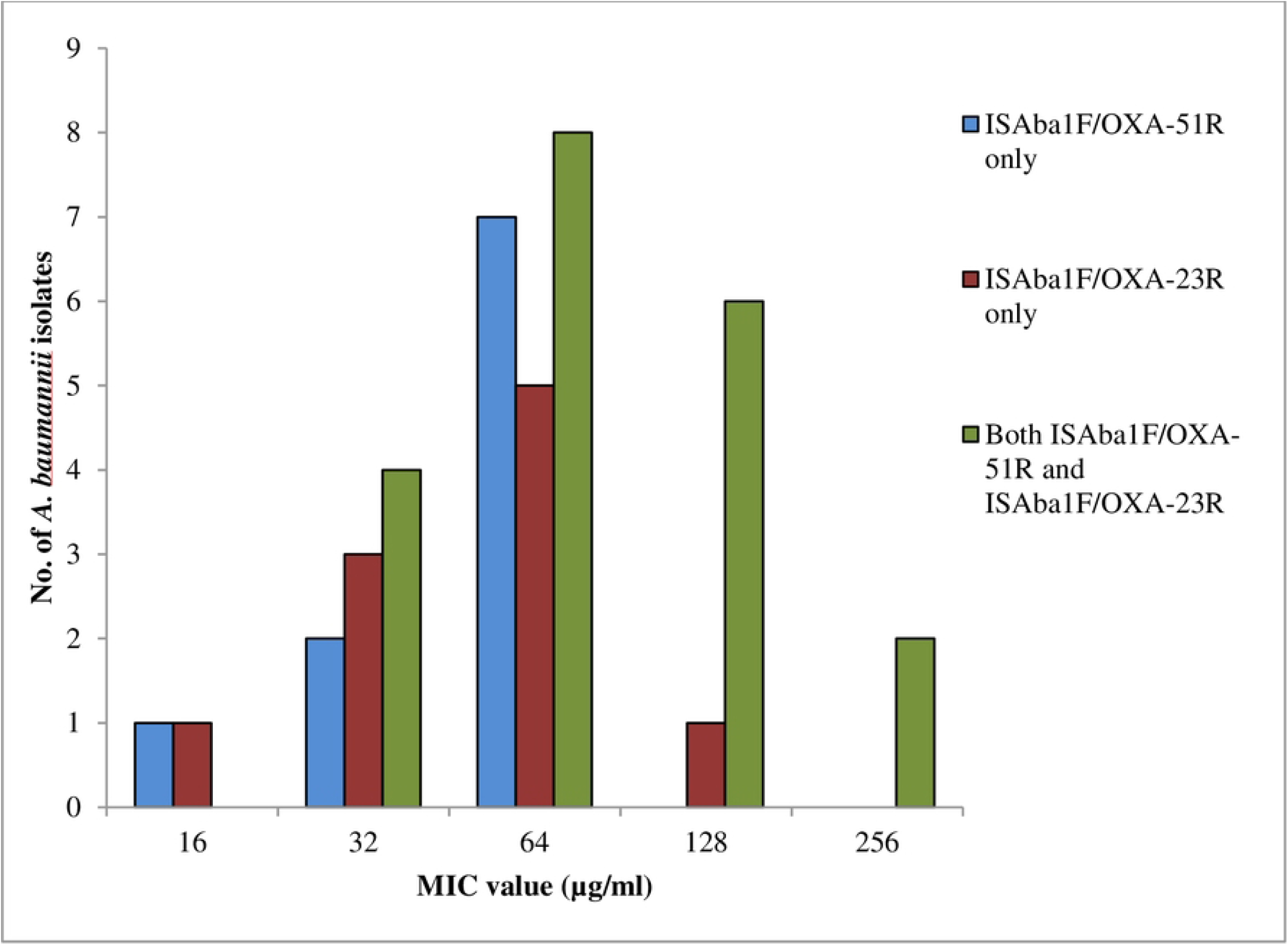
Meropenem MICs in *Acinetobacter baumannii* with *ISAba1* linked to different oxacillinase genes

## Discussion

*Acinetobacter baumannii* being notorious pathogen can cause infection in any parts of human body. *Acinetobacter baumannii* isolates were recovered from diverse clinical specimens including sputum, blood, endotracheal secretion, urine and wound swab even in this study. Most of the *A*. *baumannii* (77.5%) was isolated from samples of the respiratory tract (sputum and endotracheal secretion). This was in line with earlier investigations [13,14,27] which indicated that the respiratory system was the most often infected area by *A*. *baumannii*. Contamination of respiratory equipment is often responsible as a source of nosocomial outbreak [31].

All *Acinetobacter baumannii* isolates in this investigation were from inpatient source. This accentuates that *A*. *baumannii* has established its niche in this cardiac center. Their persistence in clinical environment is facilitated by their minimal nutrient requirements, ability to survive for long period in desiccants and resistance to key antimicrobial drugs and disinfectants [1]. Moreover, inpatients in this hospital are mostly cardiac patients which account as a high risk group for *A*. *baumannii* infection, as they are immune-compromised or have undergone some invasive procedures [32]. *Acinetobacter baumannii* has emerged to be the most prevalent nosocomial pathogen, especially in ICUs worldwide, and confers 10 – 43% mortality rate in patients admitted to ICUs [33]. Numerous studies have documented high proportion and elevated antibiotic resistance of *Acinetobacter baumannii* in ICUs [13,27,34]. Accordingly, majority of the *A*. *baumannii* isolates (60%) were recovered from ICUs in this study. Environmental contamination, particularly invasive mechanical ventilation as well as other invasive procedures frequently performed in ICUs is typically linked to nosocomial *A*. *baumannii* infection in ICUs [27].

*Acinetobacter baumannii* is increasingly being resistant to almost commonly prescribed drugs causing therapeutic failure [35]. All isolates in this research were also resistant to ciprofloxacin, piperacillin/tazobactam and extended spectrum cephalosporins. Similarly, resistance rate to gentamicin, meropenem and imipenem were also greater than 90%. These data suggested that these antibiotics are ineffective for empirical therapy against *A*. *baumannii* infection. High resistance rate was also observed against most other tested antibiotics except polymyxins and tigecycline, implying their potent usage in the treatment of *A*. *baumannii* infection in healthcare settings in Nepal. Due to their severe adverse effects and low plasma levels, these medications are typically used sparingly and only in life-threatening situations, which may explain their high susceptibility rate [36,37].

The prevalence of MDR and XDR *Acinetobacter baumannii* has risen sharply worldwide in recent years, which raises a serious concern [38]. Various studies from China, India and Iran have reported almost 100% incidence of MDR *A*. *baumannii* [2,39,40]. Similar to these studies, every single isolate of *A*. *baumannii* in this investigation were MDR and around one-third of isolates were XDR. *Acinetobacter baumannii* have high genetic plasticity with outstanding capacity to obtain various resistance determinants following lateral gene transfer, which when coupled with up-regulated innate resistance mechanism, it leads to multidrug resistance phenotype [41]. This process is even aided by the selective pressure caused by extensive misuse of antibiotics.

On phenotypic detection of β-lactamase production, 30% and 12.5% of the *Acinetobacter baumannii* were MBL and ESBL producer respectively and 2.5% isolates were KPC and AmpC producer each. This was in consistent with prior investigation [42]. But, Khanal and colleague [34] documented comparatively higher prevalence of AmpC producer (43.1%), MBL producer (54.5%) and ESBL producer (14.3%) than this study. Differences in study year and the inclusion of solely ICU patients could be the cause of the variance. Moreover, co-carriage of MBL with ESBL and KPC enzyme was also reported among *A*. *baumannii* in this study. Co-existence of various classes of β-lactamases in single bacterial isolate presents difficulties in both diagnosis and treatment [43]. There was significant association between resistance phenotypes and β-lactamase profile of *A*. *baumannii* (p=0.04) in this study. Majority of the MBL producers were XDR. The presence of integron could be associative factor between MBL and XDR phenotype. Typically, MBL genes are encoded on integron gene cassettes [44]. Along with these genes, integron also harbor multiple other antibiotic resistance gene cassettes, which results enhanced resistance to most antibiotics, leading to XDR phenotype [45]. But, further molecular study detecting integrons and other factors correlated with extensively drug resistance needs to be carried out.

All the isolates were observed to be resistant against meropenem by agar dilution method with high MIC value ranging from 16-256 μg/ml. In accordance with this study, around hundred percent resistivity to carbapenem was reported in studies from Nepal [13,14] and from adjoining country India [46], with MIC value even ranging to 512 μg/ml [14]. Nadia along with other researchers established a significant correlation between the use of carbapenem and the prevalence of resistance (r=0.778, p<0.001) [47]. As carbapenems are widely and indiscriminately prescribed in Nepal, this might elaborate the increased resistance rate to these antibiotics [48]. Resistance to carbapenems has also been associated with production of β-lactamases, modification in exterior membrane and penicillin binding proteins and overexpression of AdeABC efflux system [8].

Oxacillinase type (OXA) carbapenemases are the primary factors responsible for carbapenem resistance in *Acinetobacter baumannii* globally [49]. As phenotypic methods for determining the presence of OXA carbapenemases in *A*. *baumannii* have not yet been documented [45], PCR was used to identify different OXA carbapenemases, including *bla*_OXA-51-like_, *bla*_OXA-23-like_ and *bla*_OXA-58_. OXA-23-like enzymes are the most common oxacillinases and the primary reason behind carbapenem resistance in *A*. *baumannii* [4,30]. Thereby, the 100% resistance rate to carbapenem in this research might presumably be due to the notion that all isolates had harbored *bla*OXA-23-like gene. There are numerous reports with 100% incidence of *bla*_OXA-23-like_ gene in *A*. *baumannii* [5,13,14]. This clearly depicts inclining widespread dispersion of *bla*_OXA-23-like_ gene in *A*. *baumannii*, which could be justified by its allocation in plasmids [50]. Allied to this, IS*Aba1* insertion element is also involved in spread of *bla*OXA-23-like gene forming composite transposons [51]. Plasmids (RepAci6 and pAZJ221) and transposons (*Tn2006* and *Tn2009*) have mostly contributed on horizontal dissemination of *bla*_OXA-23-like_ gene, promoting its global distribution [4,52]. Furthermore, only 5% of isolates from this study had the *bla*OXA-58 gene. Previously it was reported only in one isolate in a study [14] from Nepal. Majorly, this gene was more common in European isolates of *A*. *baumannii* [53]. To the greatest extent of our understanding, this present study is the initial research from Nepal to document clinical isolates of *A*. *baumannii* co-harboring both IS*Aba1* and *bla*_OXA-58_ genes. The presence of such rare carbapenemases along with IS*Aba1* in this current study signals the possible increment of this gene in Nepal as *A*. *baumannii* have remarkable capacity to acquire new resistance traits thorough lateral gene transfer [41], which eventually may increase resistance to carbapenem.

Carbapenem resistance genes are primarily expressed and disseminated through the insertion sequence, IS*Aba1* in *Acinetobacter baumannii* [10]. IS*Aba1* brackets carbapenemase gene forming a transposon, which actively mobilize and thereby aids in the dissemination of carbapenemase gene [54]. Such transposons are even disseminated further through plasmids horizontally [55]. In addition, insertion sequence elements contain promoter that contributes in the overexpression of down-stream antibiotic resistance genes [56]. *Acinetobacter baumannii* isolates possess IS*Aba1* upstream of the genes *bla*_OXA-51-like_, *bla*_OXA-23-like_ and *bla*_OXA-58_, contributing to the over-production of these enzymes further facilitating resistance to carbapenem [57]. Kobs et al. [27] reported that the isolates lacking IS*Aba1* or IS*Aba1* located not near to oxacillinase genes were found to be susceptible to carbapenem. In this work, IS*Aba1* was found in every isolate either upstream of *bla*_OXA-23-like_ or *bla*_OXA-51-like_ genes. This most plausibly explains none of the isolates being susceptible to carbapenem, as IS*Aba1* aids in dissemination and overexpression of oxacillinases in these isolates. High meropenem MICs (32-256 μg/ml) with high MIC_90_ value of 243.20 μg/ml was reported among 50% of the isolates with IS*Aba1* linked upstream to both *bla*_OXA-51-like_ and *bla*_OXA-23-like_ genes. But comparatively lower MIC_90_ value of 121.60 μg/ml and 64 μg/ml was documented in isolates with IS*Aba1* upstream only of *bla*OXA-23-like and *bla*OXA-51-like genes respectively. In this respect, it can be said that increased rates of carbapenem hydrolysis could be attributed due to overexpression of both *bla*_OXA-23-like_ and *bla*_OXA-51-like_ genes by IS*Aba1*.

There were several limitations in the study. As this study was limited to single hospital, its findings cannot be generalized to the entire nation or worldwide. Confirmation of the location of insertion sequence can only be done through nucleotide sequencing, which was infeasible due to financial constraints.

In conclusion, there is agonizing state of carbapenem resistance among *Acinetobacter baumannii* in regard to high distribution of genes encoding oxacillinases and insertion sequence in this region. The emergence and spread of IS*Aba1*-linked oxacillinase genes further enhanced carbapenem resistance level creating havoc. Thus, all the medical fraternity and stakeholders need to be vigilant and enable the targeted approaches.

## Acknowledgements

We would like to thank Professor Dr. Balaji Veeraraghavan, Department of Microbiology, Christian Medical College, Vellore, India for kindly providing positive amplification controls.

